# WIO-ReefFish: A High-Resolution Dataset for Taxon-Aware Coral Reef Fish Detection in the Western Indian Ocean

**DOI:** 10.64898/2026.08.19.745797

**Authors:** Jules Gerard, Luca Branger, Filip Huyghe, Marc Kochzius, Levy Otwoma, Stephen Bergacker, Lode op’t Roodt, Cyrus Rumisha, Leandro Di Bella

**Affiliations:** Marine Biology – Ecology, Evolution & Genetics (bDIV), Vrije Universiteit Brussel (VUB), Pleinlaan 2, Brussels, Belgium; Electronics and Informatics, Vrije Universiteit Brussel (VUB), Pleinlaan 2, Brussels, Belgium; Kenya Marine and Fisheries Research Institute (KMFRI), P.O. Box 81651–80100, Mombasa, Kenya; Université Libre de Bruxelles (ULB), Avenue Franklin D. Roosevelt 50, Brussels, Belgium; Sokoine University of Agriculture (SUA), P.O. Box 3000, Chuo Kikuu, Morogoro, Tanzania

**Keywords:** Biodiversity monitoring, Coral reef fish, Diver-operated video, Object detection, Underwater computer vision, Western Indian Ocean

## Abstract

Coral reef fish assemblages are widely used as indicators of ecosystem condition, yet manual annotation of underwater video remains a major bottleneck for scalable biodiversity monitoring. Despite rapid progress in automated detection, ecologically realistic and publicly available datasets remain scarce, particularly for the Western Indian Ocean. Here, we present WIO-ReefFish, a reef fish detection dataset derived from diver-operated line-intercept transects and designed for ecological monitoring under natural survey conditions. WIO-ReefFish comprises 1,000 ultra-high-definition images (3840 × 2160 pixels) and 6,768 exhaustive bounding-box annotations spanning 24 taxonomic categories, thereby preserving full-frame assemblage structure in complex reef scenes. We also establish a standardized benchmark across nine object detection models under two complementary protocols: class-aware detection and class-agnostic fish localization. Detection performance was consistently higher under the class-agnostic protocol. The best-performing model (RT-DETR) improved from 0.48 mAP_50_ in the class-aware setting to 0.70 mAP_50_ when taxonomic constraints were removed, indicating that taxonomic discrimination remains substantially more challenging than fish localisation in reef imagery. Spatially independent evaluation revealed a pronounced generalisation gap, particularly for taxonomic detection, whereas class-agnostic fish localisation remained substantially more robust across transects and countries. Together, these results establish WIO-ReefFish as a realistic benchmark for automated reef fish detection and provide a foundation for more robust computer-vision tools in coral reef biodiversity monitoring. The WIO-ReefFish dataset and associated benchmarking resources are publicly available.

## 1. Introduction

Coral reef fish communities are widely used as indicators of ecosystem health, as they integrate habitat complexity, fishing pressure, and environmental change into measurable community-level responses [1, 2]. Along the East African coastline (EAC), reef-associated fisheries support food security and livelihoods, yet these ecosystems are increasingly threatened by climate-driven bleaching mortality and persistent over-exploitation [3–6]. Reliable monitoring of coral reef fish communities is therefore central to assessing ecological status and guiding adaptive management [7]. Traditionally, such monitoring has relied on underwater visual censuses (UVCs), which are limited in scalability and reproducibility [8]. Video-based surveys address some of these constraints by preserving permanent records and enabling standardised, quantitative analyses. A key bottleneck remains the annotation of complex reef scenes, where dense assemblages and visual ambiguity make manual labelling time-intensive and error-prone [8, 9]. Automated detection approaches offer a pathway to scale ecological monitoring, yet their effectiveness depends strongly on the datasets used to train the underlying models and evaluate their performance, particularly with respect to ecological realism. Over the past decade, fish image datasets have expanded across marine and freshwater ecosystems to support automated classification, detection, and tracking. Existing resources span a wide range of acquisition methods, including fixed underwater observatories, web-sourced image collections, controlled out-of-water imagery, and natural-habitat surveys [10–18]. While these datasets have substantially advanced automated fish image analysis, their ecological realism and suitability for coral reef monitoring vary widely. Many widely used resources are optimised for species-level classification under limited visual conditions. Fish4Knowledge, for example, was collected using fixed, relatively low-resolution cameras, limiting spatial detail for small or occluded individuals in complex scenes [10]. Several benchmarks are based on centred or cropped single-fish instances that reduce background and inter-individual complexity [19], while controlled out-of-water datasets remove key *in situ* variation in illumination, turbidity, and habitat context [14–17, 20]. Although such settings facilitate algorithm development and often yield high reported accuracy [21, 22], they differ substantially from ecological monitoring contexts, where multiple individuals coexist within dynamic and structurally complex coral reef habitats. As a result, performance on classification-oriented benchmarks does not necessarily translate to reliable full-frame detection for community-level biodiversity assessment.

Monitoring-oriented datasets derived from underwater video systems more closely reflect ecological survey conditions. These include imagery collected using Baited Remote Underwater Video Systems (BRUVS), Remote Underwater Video (RUV), and stationary observatory platforms [23–27]. Such datasets preserve full-frame scenes containing multiple individuals, but differ substantially in acquisition protocol, annotation density, taxonomic resolution, geographic scope, and intended reuse. Among publicly available resources, five existing datasets are most closely aligned with ecological monitoring objectives (Table 1).

**Table 1.** Comparative overview of image-based fish detection datasets used for ecological monitoring. Taxonomic resolution: S, species; G, genus; SF, subfamily; F, family. Survey types: RUV, remote underwater video; BRUVS, baited remote underwater video system; DOV, diver-operated video. Annotation types: bbox, bounding box; seg, segmentation mask. *Counts exclude annotations labelled only as generic “fish”.

| Name | Dataset characteristics |  |  |  |  |  | Annotations |  |  |
| --- | --- | --- | --- | --- | --- | --- | --- | --- | --- |
|  | Year | Country | Survey | Res. (px) | Taxon. | Images | *n* | Type | / img |
| Fish in Seagrass Habitat [23] | 2020 | AU | RUV | 1920×1080 | S(2) | 4,281 | 9,429 | seg | 2.2 |
| OzFish (subset) [26] | 2019 | AU | BRUV | 1920×1080 | S(12) | 1,751 | 5,112* | bbox | 2.9 |
| SEAMAPD21 [24] | 2018–2019 | US | BRUVS | 1920×1080 | S(130) | 28,328 | 90,000 | bbox | 3.2 |
| OBSEA [25] | 2013–2014 | ES | RUV | 2048×1536 | S;G (24;5) | 33,805 | 36,777* | bbox | 1.1 |
| SCSFish2025 [27] | 2017 | CN | RUV | 1920×1080 | S(28) | 11,956 | 86,934* | bbox | 7.3 |
| **WIO-ReefFish (this study)** | **2023 & 2025** | **KE/TZ** | **DOV** | **3840×2160** | **F;SF (23;1)** | **1,000** | **6,768** | **bbox** | **6.8** |

The datasets listed in Table 1 differ primarily in their acquisition paradigms, which influence scene composition, fish behaviour, and background dynamics. Fish in Seagrass Habitat [23], OzFish [26], and SEAMAPD21 [24] rely on remote or baited underwater video systems, enabling standardised sampling across large spatial extents and multiple habitats through repeated camera placements. Although such deployments may be repeated across sites to increase spatial coverage, recordings are characterised by fixed viewpoints and relatively static backgrounds within individual videos. In the case of BRUVS, bait can further influence observed assemblages by preferentially attracting predators and scavengers, thereby biasing local species composition and relative abundance estimates [28–30]. Consequently, although BRUVS are effective for detecting spatial patterns and increasing encounter rates, the resulting community representation may not fully reflect natural assemblages. A distinct stationary paradigm is represented by fixed underwater observatory cameras, as exemplified by OBSEA [25] and SCSFish2025 [27]. Such systems can support frequent observations and dense, species-level annotation. However, their fixed viewpoints produce consistent backgrounds and site-specific visual conditions that may constrain generalisation to broader surveys. In contrast, diver-operated video (DOV) transects, which are widely used in coral reef ecology, record continuous habitat gradients and dynamic backgrounds along survey paths [31]. These conditions introduce additional visual complexity through occlusion, turbidity, heterogeneous substrates, camera movement, and changing viewing angles [20, 32, 33]. Although DOV transects may underrepresent cryptic or diver-avoidant species, they capture spatial variation in substrate, visibility, and fish occurrence across coral reef habitats. Unlike baited systems, diver-operated transects do not deliberately attract individuals and therefore provide a complementary representation of fish assemblages under standard transect-survey conditions.

Existing monitoring-oriented datasets also differ fundamentally in how completely and consistently fish assemblages are represented within frames. In Fish in Seagrass Habitat [23], instance-level segmentation is applied to a small number of target species, consistent with a focal-species monitoring objective rather than comprehensive community annotation. OzFish [26], SEAMAPD21 [24], OBSEA [25], and SCSFish2025 [27] provide frame-level annotations that encompass broader portions of the assemblage, although not all individuals are assigned to the same taxonomic resolution. Some individuals are instead retained under generic, mixed, or unclassified taxonomic categories because of annotation constraints and visual ambiguity. Annotation density further conditions how these strategies operate in practice. Datasets derived from structurally complex habitats may contain dense assemblages with frequent occlusion and inter-individual overlap, which are relevant to biodiversity monitoring but substantially increase annotation difficulty. OBSEA [25] and SCSFish2025 [27] both include crowded, multi-individual scenes recorded by fixed cameras, resulting in limited background variation within individual videos. Mobile surveys can additionally combine high annotation density with camera-induced motion and continuously changing habitat structure. Image resolution further differentiates existing datasets and influences both annotation quality and detection performance. Most monitoring-oriented datasets rely on high-definition imagery with resolutions up to 1920×1080 [23, 24, 26, 27] or 2048×1536 pixels [25]. This can constrain the representation of small or distant individuals in visually complex scenes. Ultra-high-definition imagery preserves finer spatial detail and may support more complete and precise annotation, particularly in mobile transect surveys where fish occur across a wide range of sizes, distances, and viewing angles. These dataset limitations are particularly relevant in the Western Indian Ocean (WIO), a major coral biodiversity hotspot that ranks among the world’s most species-rich reef regions after the Coral Triangle [34, 35]. Publicly accessible fish-detection datasets remain particularly limited in the region [33, 36]. Previous automated fish-classification studies in the region have relied primarily on private datasets [22], constraining reproducibility, methodological comparison, and reuse.

Together, these limitations indicate that existing datasets and benchmarks do not fully capture the ecological and observational complexity inherent to reef-monitoring surveys. In particular, there remains a need for a publicly accessible dataset that combines high spatial resolution, exhaustive full-frame annotations, consistent taxonomic treatment, and regional representation in the WIO. Here, we present WIO-ReefFish, a reef-fish detection dataset derived from DOV transects in Kenya and Tanzania. All sufficiently visible and identifiable fish were annotated using a consistent coarse taxonomic scheme comprising 24 taxa. The dataset retains full-frame scenes with multiple co-occurring fish under realistic reef survey conditions and remains extensible to finer taxonomic resolution without modification of the underlying spatial annotations. WIO-ReefFish is accompanied by a benchmark of representative object-detection architectures under class-aware and class-agnostic evaluation. Together, the dataset and benchmark provide a reference point for evaluating automated reef-fish detection under ecologically realistic survey conditions.

Our principal contributions are:

- **WIO-ReefFish dataset.** We introduce WIO-ReefFish, to our knowledge the first publicly available reef-fish detection dataset from the WIO derived from DOV surveys. It provides ultra-high-definition imagery with exhaustive full-frame annotations across 23 families and one subfamily, derived from video transects spanning multiple coral reef sites in Kenya and Tanzania.
- **Detection benchmark.** We establish a standardised evaluation benchmark for WIO-ReefFish, comparing multiple object-detection architectures under two complementary task definitions: class-aware detection and class-agnostic fish localisation.

## 2. WIO-ReefFish Dataset

The WIO-ReefFish dataset was developed to support automated detection of coral reef fish under realistic survey conditions. Representative frames are shown in Fig. 1. This section outlines its design, acquisition, annotation protocol, and composition.

**Figure 1:**
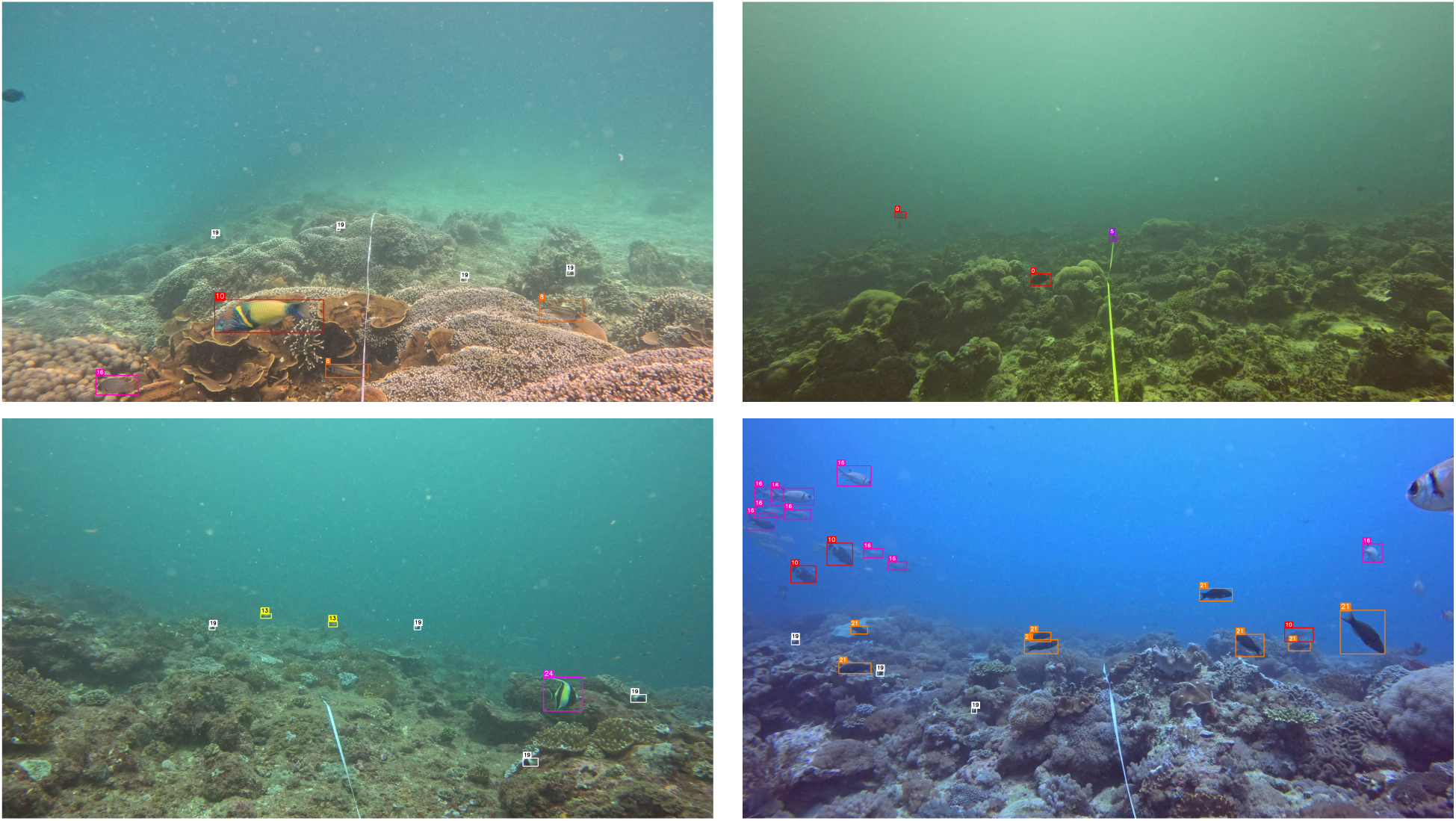
Representative frames from the WIO-ReefFish dataset illustrating variation in reef scenes and fish assemblages across sampling sites along the EAC.

### 2.1. Design and taxonomic scope

Fish meeting the annotation criteria were assigned using a consistent coarse taxonomic scheme comprising 23 families and one subfamily (Scarinae). Scarinae was retained as a separate category because it was morphologically distinguishable from the remaining Labridae and represents a functionally distinct reef fish group. This taxonomic coverage reflects the sampled imagery and does not encompass all reef-fish families occurring in the WIO. The selected resolution balances taxonomic consistency with the practical constraints of underwater imagery, where species-level identification is frequently limited by occlusion, visibility, and viewing angle. Coarse taxonomic resolution nevertheless retains ecologically relevant functional information because reef-fish trophic guilds are strongly phylogenetically conserved [37]. The spatial annotations remain extensible to finer taxonomic resolution in future dataset versions.

### 2.2. Data acquisition

DOV transects were conducted during two field campaigns along the EAC. Surveys in March 2023 targeted reef sites along the Kenyan coastline and Zanzibar (Tanzania), while a second campaign in February 2025 expanded coverage along the Tanzanian mainland and offshore islands. Surveys covered 17 reef sites, including nine in Kenya and eight in Tanzania, distributed among 10 coastal locations and spanning a range of habitat conditions and management regimes (Fig. 2; Appendix A).

**Figure 2:**
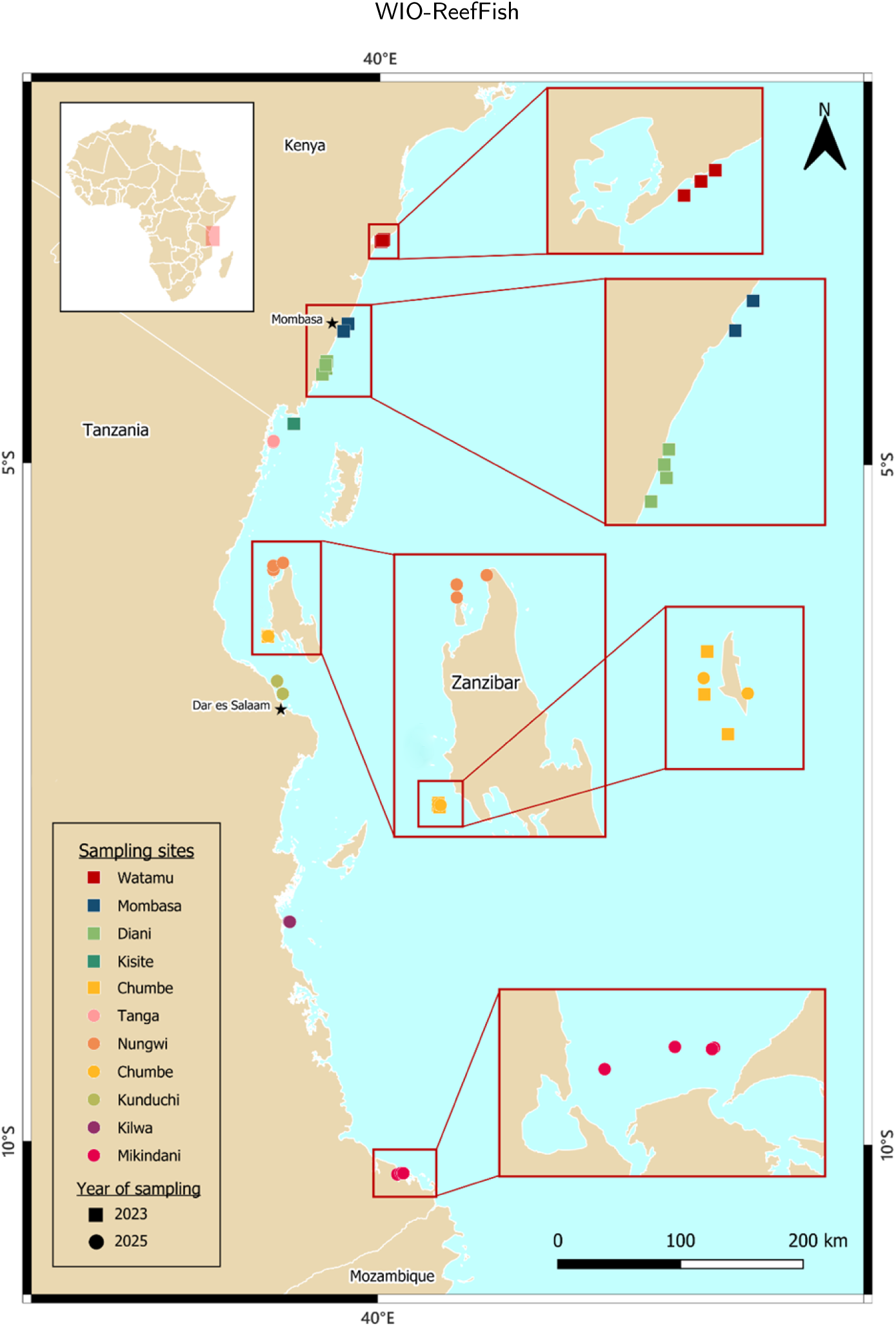
Sampling locations of WIO-ReefFish DOV transects along the Kenyan and Tanzanian coast.

DOV transects followed a standardised protocol [31]. At each site, 50 m transects were deployed at depths of 3–16 m and recorded using forward-facing action cameras (GoPro Hero 11 Black) positioned approximately 1 m above the reef. After a five minute settling period to compensate for possible fish disturbance during transect deployment, transects were filmed at a constant pace (approximately 3 minutes per transect). Between four and ten transects were conducted per site, yielding a total of 81 video sequences (approximately 4.1 hours of footage). Video frames were extracted at regular temporal intervals (every 3 s). After frame extraction and pre-screening for visible fish presence, 47 of these 81 transects contributed at least one annotated frame to the final dataset; the remaining 34 yielded no frames meeting the inclusion criteria.

### 2.3. Annotation protocol and quality control

Fish were manually annotated following a consistent, literature-informed protocol for underwater object detection that ensures sufficient visual and spatial quality [38, 39]. Bounding boxes were drawn tightly around the visible body silhouette. Partially occluded individuals were retained when diagnostic features remained discernible and at least 50% of the body was visible. A minimum bounding-box size of 500 px^2^ was applied to ensure sufficient spatial information for reliable localisation and classification while retaining small-bodied taxa relevant to biodiversity monitoring. Individuals cropped at the image boundary, severely affected by blur or turbidity, or exhibiting ambiguous morphology were excluded when taxonomic identification could not be reliably established. Annotation criteria prioritised views that preserved diagnostic features (e.g. lateral or oblique orientations), ensuring consistency and taxonomic accuracy across the dataset.

All fish meeting these predefined criteria were then identified using the *Coastal Fishes of the Western Indian Ocean* guide [40], which resulted in a coarse taxonomic scheme comprising 24 taxa. Lastly, identifications were independently reviewed by experienced reef fish taxonomists, with ambiguous cases resolved by consensus.

### 2.4. Dataset composition

The final dataset comprises 1,000 images containing 6,768 annotated fish instances across 24 taxonomic categories, including 23 families and one subfamily. Kenyan and Tanzanian coral reefs contributed equally to the image dataset. Image availability averaged 98.8 images per location (range 9–171) and 58.1 images per site (range 9–139). The class distribution is strongly long-tailed (Fig. 3), with damselfish (Pomacentridae) accounting for approximately 55% of all annotations, while 15 taxa each contribute fewer than 1% of instances.

**Figure 3:**
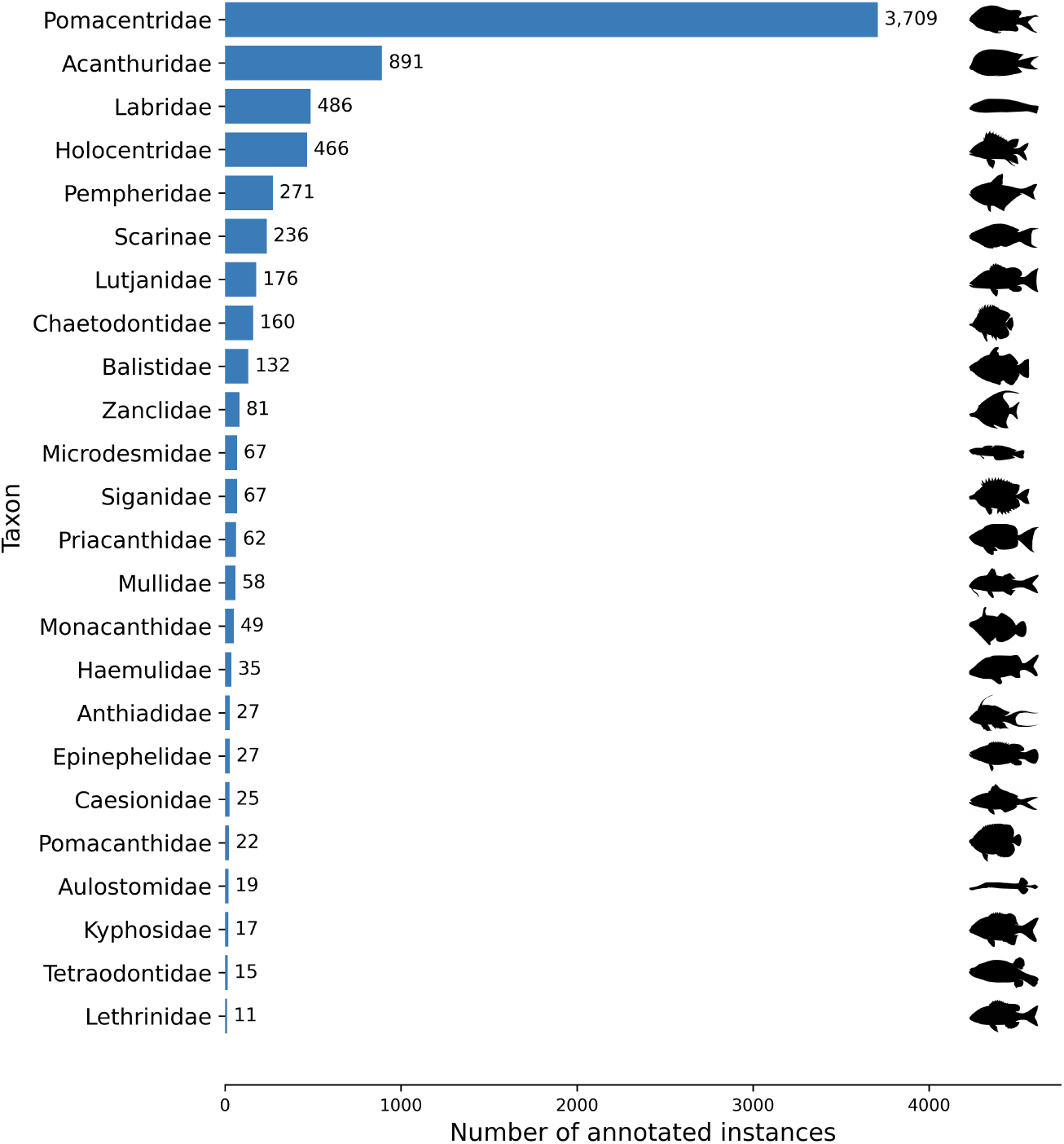
Distribution of annotated fish taxa instances in the WIO-ReefFish dataset. Representative silhouettes are not shown to scale.

At the image level, annotation density and taxonomic composition vary substantially (Fig. 4), with an average of 6.8 instances per image (range 1–84) and approximately 2.4 taxa per image. Assemblages are frequently dominated by a single class (mean dominance = 71.7%), while bounding boxes are predominantly small relative to the full image area (median = 0.06%). All images are provided at ultra-high-definition resolution (3840×2160 pixels).

**Figure 4:**
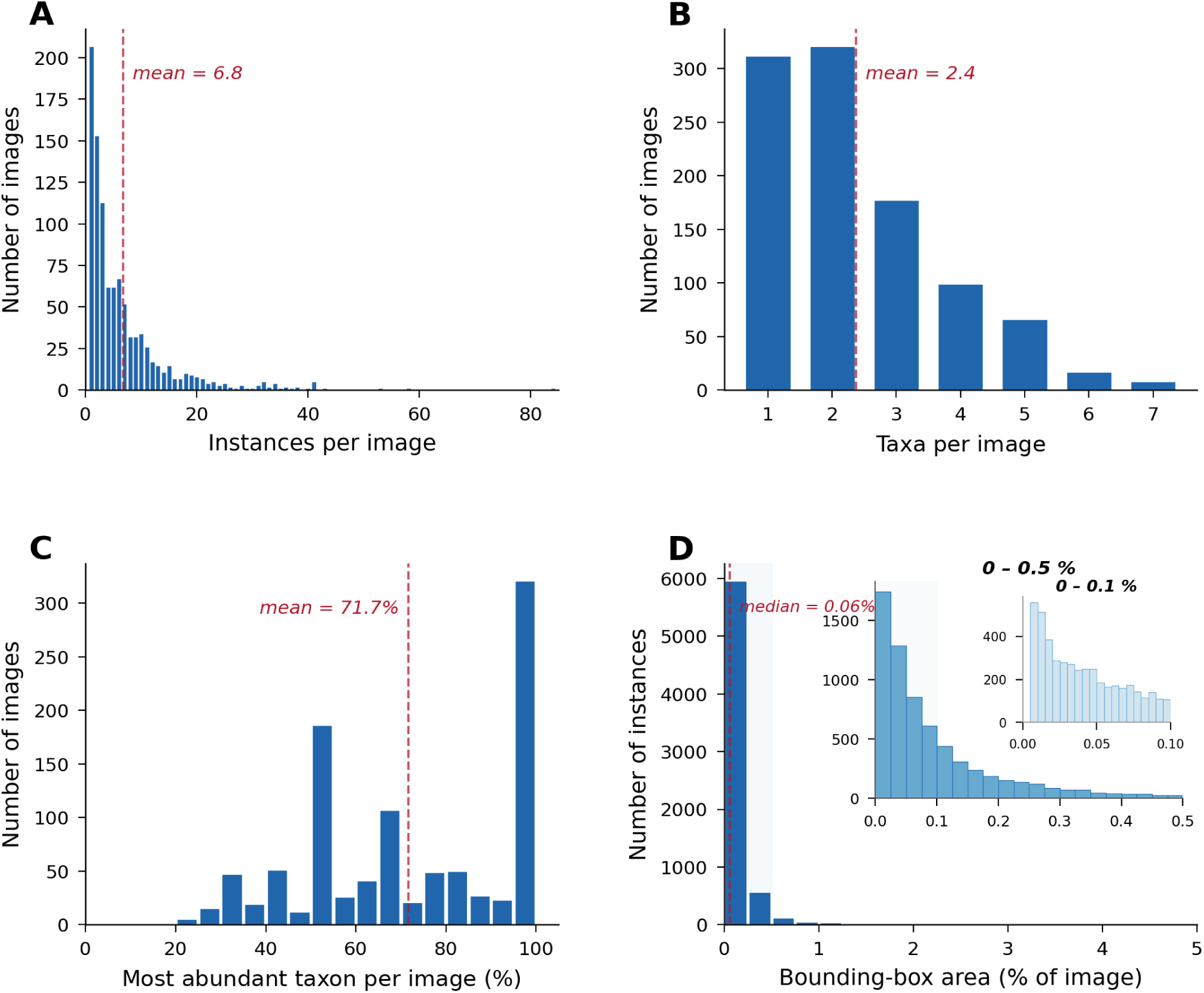
Image- and instance-level characteristics of the WIO-ReefFish dataset obtained from DOV transects conducted in Kenyan and Tanzanian coral reefs. (A) Number of annotated instances per image. (B) Number of taxonomic categories per image. (C) Image-level dominance, defined as the proportion of individuals belonging to the most abundant taxon, indicating that many images are strongly dominated by a single taxon. (D) Relative bounding-box size expressed as a percentage of image area, with nested insets highlighting the prevalence of small and very small objects. Together, these distributions illustrate the high object density, taxonomic imbalance, and small-object dominance that characterise the dataset.

## 3. Experimental Setup

To provide a reference point for future studies, WIO-ReefFish was benchmarked using nine representative object-detection architectures, including one-stage detectors such as You Only Look Once (YOLO) [41], two-stage models such as Faster Region-based Convolutional Neural Network (Faster R-CNN) [42], and transformer-based approaches such as Real-Time Detection Transformer (RT-DETR) [43]. The benchmark used the full dataset of 1,000 images and 6,768 bounding-box annotations across 24 taxonomic categories. Images were divided into 700 training, 150 validation, and 150 test images, and all models were initially evaluated at an input resolution of 640 × 640 pixels.

### 3.1. Evaluation protocols

Two complementary protocols are applied:

- **Class-aware detection (24 taxonomic categories)**: A predicted box was considered a true positive when it overlapped a ground-truth box by at least IoU = 0.50 and the predicted taxonomic category matched the corresponding ground-truth label. Intersection over union (IoU) is the area of overlap between the predicted and ground-truth boxes divided by the area of their union. This protocol measures both localisation and taxonomic classification.
- **Class-agnostic localisation (fish vs. background):** All taxonomic labels were remapped to a single “fish” category before evaluation. A predicted box was considered a true positive when it overlapped any ground-truth fish box by at least IoU = 0.50, irrespective of taxonomy. This protocol isolates fish localisation and enables comparison with zero-shot models such as Grounding DINO, which return only a generic “fish” category.

### 3.2. Metrics

We define a standardised evaluation protocol for object detection on reef imagery. We report precision (*P*), recall (*R*), F1 score, mean average precision at IoU = 0.50 (mAP_50_), and COCO-style mAP averaged over IoU thresholds from 0.50 to 0.95 in increments of 0.05 (mAP_50-95_). Precision is the proportion of predicted detections that are correct, recall is the proportion of ground-truth fish instances that are detected, and F1 is their harmonic mean. mAP_50_ was computed using PASCAL VOC 11-point interpolation:

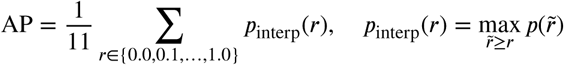

where *p*(*r̃*) is the precision at recall *r̃*. For the class-aware protocol, mAP_50_ and mAP_50-95_ were averaged across all 24 classes.

### 3.3. Difficulty levels

Three difficulty levels were defined from the distribution of ground-truth bounding-box heights. The **Easy** level evaluates large objects only, the **Moderate** level includes medium and large objects, and the **Hard** level includes all annotated objects. Ground-truth boxes below the threshold associated with each level were marked as *ignored* during evaluation. The resulting thresholds were 64 pixels for Easy and 32 pixels for Moderate, yielding approximate Easy/Moderate/Hard partitions of 46%, 34%, and 20% of annotations, respectively. Full definitions are provided in Appendix Table A.2.

### 3.4. Training configuration

All supervised baselines were trained for 100 epochs with early stopping (patience = 20 epochs) using an input resolution of 640 × 640 pixels and batch size 16 (8 for the 1280 × 1280 ablation). Optimisation was performed using AdamW or SGD depending on the detector architecture. Standard geometric and photometric augmentations were applied following Ultralytics defaults. To assess training variability, RT-DETR, YOLO11, and YOLO26 were each trained using three independent random seeds (0, 1, and 2). Unless otherwise stated, results for these models are reported as mean±standard deviation across the three runs, whereas results for all other models are based on a single run. All experiments were conducted on a single NVIDIA RTX 4090 GPU. Additional implementation details are provided in Appendix B.

Grounding DINO was evaluated in zero-shot mode using prompts constructed from the WIO-ReefFish taxonomic categories, whereas YOLO-World was fine-tuned on the WIO-ReefFish training split using the same category prompts.

DINOv2 (ViT-S/14) was evaluated using a lightweight anchor-free detection head trained on frozen backbone features.

## 4. Results

### 4.1. Overall benchmark performance

Overall detection performance under both evaluation protocols is reported in Table 2. The top three baselines (RT-DETR, YOLO11, YOLO26) were each averaged across three independent training seeds. RT-DETR achieved the highest mAP_50_ under both class-aware and class-agnostic evaluation, reaching 0.480±0.015 and 0.698±0.009 mAP_50_, respectively. In contrast, the YOLO-based detectors consistently produced higher precision and F1 scores, with YOLO11 achieving the highest precision under both protocols (0.602 class-aware, 0.683 class-agnostic) and the highest class-agnostic F1 (0.664), while YOLO26 led on class-aware F1 (0.592). All models performed substantially better under the class-agnostic protocol, indicating that fish localisation was markedly easier than taxonomic classification. Across architectures, the persistent gap between mAP_50_ and mAP_50-95_ further indicates reduced localisation accuracy at stricter IoU thresholds.

**Table 2.**
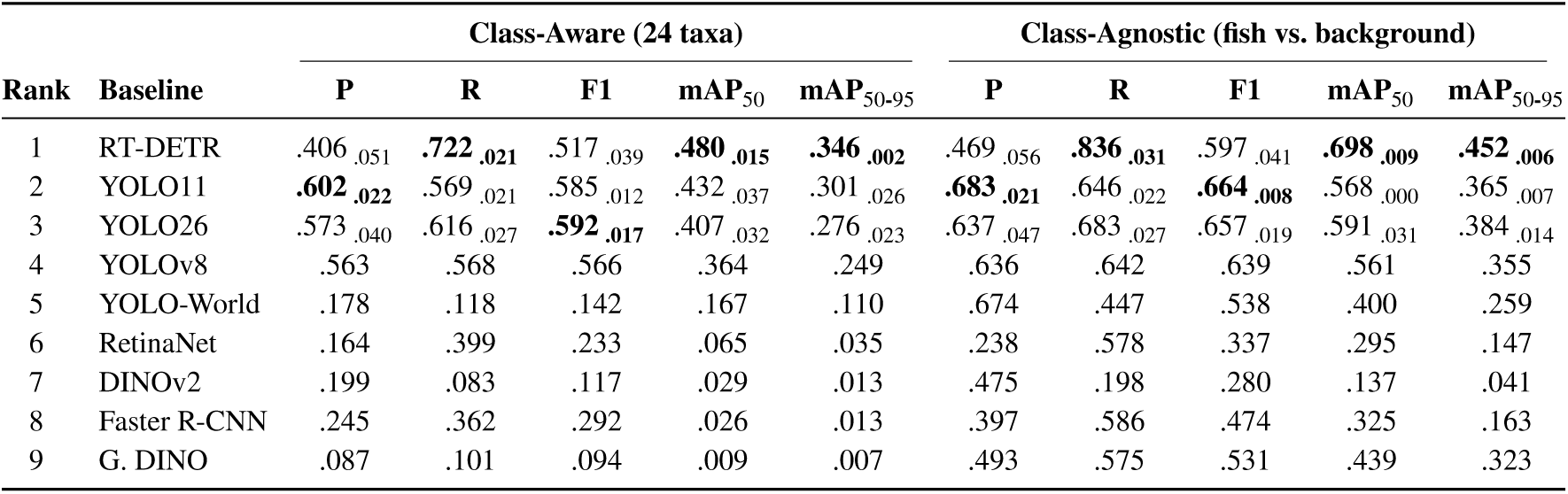
Overall detection performance on the test split under both evaluation protocols, ranked by class-aware mAP_50_. Subscripted values denote standard deviations.

| Rank | Baseline | Class-Aware (24 taxa) |  |  |  |  | Class-Agnostic (fish vs. background) |  |  |  |  |
| --- | --- | --- | --- | --- | --- | --- | --- | --- | --- | --- | --- |
| | | P | R | F1 | $mAP_{50}$ | $mAP_{50-95}$ | P | R | F1 | $mAP_{50}$ | $mAP_{50-95}$ |
| 1 | RT-DETR | .406<sub>.051</sub> | **.722**<sub>.021</sub> | .517<sub>.039</sub> | **.480**<sub>.015</sub> | **.346**<sub>.002</sub> | .469<sub>.056</sub> | **.836**<sub>.031</sub> | .597<sub>.041</sub> | **.698**<sub>.009</sub> | **.452**<sub>.006</sub> |
| 2 | YOLO11 | **.602**<sub>.022</sub> | .569<sub>.021</sub> | .585<sub>.012</sub> | .432<sub>.037</sub> | .301<sub>.026</sub> | **.683**<sub>.021</sub> | .646<sub>.022</sub> | **.664**<sub>.008</sub> | .568<sub>.000</sub> | .365<sub>.007</sub> |
| 3 | YOLO26 | .573<sub>.040</sub> | .616<sub>.027</sub> | **.592**<sub>.017</sub> | .407<sub>.032</sub> | .276<sub>.023</sub> | .637<sub>.047</sub> | .683<sub>.027</sub> | .657<sub>.019</sub> | .591<sub>.031</sub> | .384<sub>.014</sub> |
| 4 | YOLOv8 | .563 | .568 | .566 | .364 | .249 | .636 | .642 | .639 | .561 | .355 |
| 5 | YOLO-World | .178 | .118 | .142 | .167 | .110 | .674 | .447 | .538 | .400 | .259 |
| 6 | RetinaNet | .164 | .399 | .233 | .065 | .035 | .238 | .578 | .337 | .295 | .147 |
| 7 | DINOv2 | .199 | .083 | .117 | .029 | .013 | .475 | .198 | .280 | .137 | .041 |
| 8 | Faster R-CNN | .245 | .362 | .292 | .026 | .013 | .397 | .586 | .474 | .325 | .163 |
| 9 | G. DINO | .087 | .101 | .094 | .009 | .007 | .493 | .575 | .531 | .439 | .323 |

### 4.2. Taxonomic performance

Performance on the full 24 class benchmark may be influenced by the long-tailed class distribution, where several fish taxa are represented by very few training instances. To assess the contribution of low-support taxa, the class-aware evaluation was repeated, using only the eight most frequent taxa (≥ 100 training instances), representing 91% of all test annotations (Table 3).

**Table 3.** Class-aware results restricted to the eight most frequent taxa (≥ 100 training instances; 91% of test annotations), with full-set results reproduced for direct comparison. Subscripted values denote standard deviations.

| Baseline | $mAP_{50}$ | | $mAP_{50-95}$ | |
| --- | --- | --- | --- | --- |
|  | Freq. | All | Freq. | All |
| RT-DETR | **.540**<sub>.011</sub> | **.480**<sub>.015</sub> | **.380**<sub>.004</sub> | **.346**<sub>.002</sub> |
| YOLO11 | .448<sub>.002</sub> | .432<sub>.037</sub> | .310<sub>.003</sub> | .301<sub>.026</sub> |
| YOLO26 | .462<sub>.011</sub> | .407<sub>.032</sub> | .319<sub>.009</sub> | .276<sub>.023</sub> |
| YOLOv8 | .433 | .364 | .299 | .258 |
| YOLO-World | .139 | .167 | .100 | .120 |

Restricting evaluation to frequent taxa confirmed RT-DETR as the best-performing detector. Class-aware mAP_50_ values increased substantially for all top supervised detectors when low-support taxa were excluded, with RT-DETR reaching 0.540 and the YOLO family clustering around 0.45.

The RT-DETR confusion matrix (Fig. C.2) shows that taxonomic misclassifications were concentrated among dominant and intermediate-frequency taxa, whereas rare taxa exhibited sparse and unstable detection patterns. Background-related errors remained common, and included both missed detections and false positives. For comparison, the corresponding confusion matrix for YOLO11 is provided in Fig. C.3.

Per-class performance varied substantially and did not follow class frequency alone. Across-seed variability was small for well-represented taxa (e.g. Pomacentridae: AP_50_ standard deviation = 0.014) but substantial for low-support taxa (e.g. Zanclidae: 0.138, with only 12 test instances). Several well-represented taxa remained difficult despite relatively high support, whereas some visually distinctive taxa achieved high AP values with comparatively few examples. Full per-class metrics are reported in Appendix Table C.1.

### 4.3. Geographic generalisation

The benchmark above uses a random image-level train/validation/test split. However, an audit revealed substantial within-transect overlap between subsets, with 41 of 47 transect videos contributing frames to all three splits. To evaluate generalisation more rigorously, we constructed two additional partitioning strategies and re-trained the five strongest supervised baselines under identical training conditions.

The first (data_split_3) was a transect-stratified split in which each video was assigned exclusively to either train, validation or test while maintaining geographic representation across locations. The second (data_split_4) used country-level holdout evaluation, training on one country and testing on the other (KE→TZ and TZ→KE), to assess geographic transfer across distinct survey domains. Random-split results from Table 2 are reproduced for comparison.

Three main patterns emerged. First, class-agnostic localisation remained relatively robust under stricter split schemes (Table 4). RT-DETR decreased only modestly between the random and transect-stratified splits (0.698 to 0.635 mAP_50_). Second, taxonomic classification was substantially more sensitive to splitting strategy. Under the transect-stratified split, class-aware mAP_50_ decreased sharply across all models, with RT-DETR dropping from 0.480 to 0.230. YOLO-based detectors similarly converged toward substantially lower performance levels (0.185–0.188). Third, country-level transfer introduced an additional but smaller performance decrease relative to the transect-stratified split. Relative to the transect-stratified split, RT-DETR’s class-agnostic mAP_50_ decreased by a further 0.061 under KE→TZ and by 0.033 under TZ→KE.

**Table 4.** Class-agnostic and class-aware mAP_50_ under four train/test partitioning strategies. *Random* corresponds to the original image-level split; *Transect* assigns each video exclusively to train, validation, or test; KE→TZ and TZ→KE denote country-level holdout evaluation.

| Baseline | Class-Agnostic $mAP_{50}$ | | | | Class-Aware $mAP_{50}$ | | | |
| --- | --- | --- | --- | --- | --- | --- | --- | --- |
|  | Rand. | Trans. | KE→TZ | TZ→KE | Rand. | Trans. | KE→TZ | TZ→KE |
| RT-DETR | **.698** | **.635** | **.574** | **.602** | **.480** | **.230** | .147 | **.153** |
| YOLO11 | .568 | .545 | .458 | .409 | .386 | .188 | .130 | .114 |
| YOLO26 | .562 | .495 | .468 | .398 | .363 | .188 | **.147** | .112 |
| YOLOv8 | .561 | .488 | .463 | .379 | .364 | .185 | .118 | .082 |
| YOLO-World | .400 | .410 | .319 | .324 | .167 | .047 | .031 | .025 |

Per-location performance was evaluated for RT-DETR under the class-agnostic protocol across the 10 sampled coastal locations in Kenya and Tanzania (Appendix Table D.1). Performance varied substantially among locations (*F̄*_1_ = 0.643 ± 0.149). The strongest and weakest scores occurred at low-sample locations, with Kunduchi reaching F1 = 1.00 (*n*=2) and Kilwa reaching F1 = 0.387 (*n*=6). Locations with larger test sets showed more intermediate performance values, including Tanga (0.649; *n*=22), Watamu (0.664; *n*=27) and Kisite (0.542; *n*=18).

### 4.4. Effect of input resolution

All preceding experiments used an input resolution of 640 × 640 pixels. To assess whether higher-resolution inference improved performance on ultra-high-definition WIO-ReefFish imagery (3840×2160), the five strongest supervised baselines were re-trained and evaluated at 1280 × 1280 resolution under both evaluation protocols.

Increasing input resolution consistently improved performance across all models (Table 5). Under class-aware evaluation, RT-DETR remained the highest-performing detector, reaching 0.614 mAP_50_ (+0.134 vs. the 640 × 640 headline), while the YOLO family showed gains of +0.049 to +0.146. Similar improvements were observed under class-agnostic evaluation, with RT-DETR reaching 0.756 mAP_50_ and YOLO11 leading on F1 (0.756).

**Table 5.** Detection performance at 1280 × 1280 input resolution on the test split under both evaluation protocols. Δ indicates the change in mAP_50_ relative to the corresponding 640 × 640 result reported in Table 2.

| Rank | Baseline | Class-Aware |  |  |  |  | Class-Agnostic |  |  |  |  |
| --- | --- | --- | --- | --- | --- | --- | --- | --- | --- | --- | --- |
| | | P | R | F1 | $\text{mAP}_{50}$ | $\Delta$ | P | R | F1 | $\text{mAP}_{50}$ | $\Delta$ |
| 1 | RT-DETR | .587 | **.804** | .679 | **.614** | +0.134 | .639 | **.875** | .739 | **.756** | +0.058 |
| 2 | YOLO11 | **.727** | .679 | .702 | .500 | +0.068 | **.783** | .731 | **.756** | .680 | +0.112 |
| 3 | YOLO26 | .709 | .712 | **.711** | .456 | +0.049 | .747 | .751 | .749 | .671 | +0.080 |
| 4 | YOLOv8 | .638 | .736 | .683 | .475 | +0.111 | .688 | .794 | .737 | .672 | +0.111 |
| 5 | YOLO-World | .186 | .198 | .192 | .313 | +0.146 | .650 | .693 | .671 | .586 | +0.186 |

### 4.5. Stratified performance analyses

Stratified analyses were performed to evaluate detector performance under varying image conditions using a fixed deployment operating point (*τ* = 0.25, IoU ≥ 0.50). Unlike the main benchmark, which reported mAP_50_ across confidence thresholds, these analyses used F1 score at the fixed operating threshold.

Performance across object-size difficulty levels is summarised in Appendix Table D.2. Most detectors showed only moderate degradation from Easy to Hard conditions, and overall ranking patterns remained stable. RT-DETR achieved the highest mAP_50_ across all difficulty levels under both evaluation protocols. Under class-aware evaluation, YOLO11 and YOLOv8 showed the smallest decreases from Easy to Hard (−0.031 and −0.030, respectively), while RT-DETR decreased by 0.046. Under class-agnostic evaluation, the strongest decline was observed for Grounding DINO (−0.178), whereas YOLO26 remained comparatively stable (−0.004).

Additional analyses of brightness-stratified performance, ensemble fusion and inference-time image enhancement are provided in Appendix D.

## 5. Discussion

WIO-ReefFish demonstrates that DOV transect surveys can serve as a viable foundation for training and benchmarking automated reef-fish detection systems. By combining full-frame DOV imagery with consistent coarse taxonomic annotations across multiple reef sites, WIO-ReefFish captures the visual and taxonomic complexity characteristic of transect-based ecological surveys.

Across all nine architectures, class-agnostic localisation consistently outperformed class-aware detection (Table 2), showing that fish localisation was more reliable than correct per-instance taxonomic assignment.

### 5.1. Taxonomic performance and long-tail structure

The long-tailed class distribution was reflected in a consistent bias toward dominant taxa: ambiguous fish detections were frequently assigned to Pomacentridae, thereby potentially inflating its apparent abundance while suppressing estimates for rarer taxa. This bias does not eliminate errors within Pomacentridae itself, but instead redistributes misclassification toward dominant taxa. Fifteen of the 24 taxa account for fewer than 1% of annotations, making their per-class estimates unstable. Restricting evaluation to the eight taxa with at least 100 training instances increased RT-DETR mAP_50_ from 0.480 to 0.540, confirming that low-support taxa depress the macro-averaged score, although substantial errors remained among well-represented taxa (Table 3).

A related pattern, evident in the confusion matrices (Figs. C.2 and C.3), is the prevalence of background-related errors in both directions: true fish instances are frequently missed, while background regions generate false-positive fish detections, often assigned to Pomacentridae. This bidirectional error pathway forms a fundamental detection uncertainty in structurally complex reef environments that is largely independent of taxonomic classification and is conserved across model architectures. Per-class performance provides additional nuance (Appendix Table C.1). Labridae and Scarinae were among the most challenging taxa despite relatively high training support. This is notable because Scarinae was separated from the remaining Labridae to form a more morphologically and functionally coherent category, yet both classes remained difficult. Their performance therefore suggests that taxonomic subdivision alone does not necessarily produce visually separable detection classes. Conversely, several morphologically distinctive taxa achieved high per-class mAP_50_ despite lower support, indicating that visual separability influences class-aware performance in addition to class frequency.

The two highest-performing models expressed this trade-off differently: RT-DETR favoured recall at the cost of more false positives, whereas YOLO11 achieved higher precision but missed more fish. Model choice therefore depends on whether the monitoring application prioritises avoiding missed individuals or limiting false abundance estimates.

### 5.2. Spatial generalisation and site-level variability

Detection performance varied substantially among the sampled locations, although estimates for locations with very small test sets were unstable (Appendix Table D.1). Descriptive associations between location-level F1 and background luminance or contrast were weak, suggesting that optical conditions alone were insufficient to explain the observed heterogeneity. Given the small number of locations, however, these correlations should be treated as exploratory.

Class-agnostic localisation remained comparatively robust under stricter splits, whereas class-aware detection declined markedly when transects were separated between training and testing (Table 4). The much larger decline under class-aware evaluation indicates that correct taxonomic assignment generalised less effectively than fish localisation. The contrast between random image-level and transect-stratified performance demonstrates that distributing frames from the same transects across training and test sets can substantially inflate apparent class-aware performance. Country-level holdout produced a smaller additional reduction than transect separation in the present experiments, suggesting that across-transect variation was the dominant generalisation challenge.

### 5.3. Future directions and operational implications

Detection performance generally declined as progressively smaller objects were included, while increasing input resolution improved all evaluated detectors (Appendix Table D.2; Table 5). Higher resolution may therefore benefit applications requiring small-object or class-aware detection, although these gains must be balanced against computational cost and processing throughput. Weighted box fusion also improved performance, whereas inference-only image enhancement reduced F1 (Appendix D), indicating that model and ensemble optimisation were more effective than post-hoc enhancement.

Further development should prioritise both dataset diversity and taxonomic design. Additional imagery should expand the representation of fish appearances, sizes and viewing angles, as well as backgrounds, habitats and optical conditions, while targeted sampling of underrepresented and frequently confused taxa would improve class balance and estimate stability. A larger resource would also permit task-specific subsets, although rebalanced benchmarks should remain distinct from datasets intended to preserve natural long-tailed assemblage structure.

Future extensions should move beyond independent frame-level bounding boxes. Multi-object tracking could reduce repeated counts, support temporal analyses, and integrate multiple views of the same fish. Segmentation may improve object delineation and shape representation, whereas calibrated stereo-video is better suited to reliable length and biomass estimation. Combined with broader spatial coverage, these developments could support repeatable indicators of fish occurrence, relative abundance, community composition, trophic structure, and biomass for reef monitoring, marine protected area assessment, and fisheries management.

## 6. Conclusion

In this study, we present WIO-ReefFish, a 4K annotated dataset for reef fish detection in the WIO, comprising 1,000 images and providing a benchmark for evaluating state-of-the-art object detection models under realistic reef survey conditions. Across all evaluated architectures, fish localisation consistently proved more reliable than taxonomic classification, highlighting the difficulty of assigning the correct taxonomic category under natural reef conditions. Additional analyses further showed strong effects of long-tailed taxonomic structure, ecological variability across reef sites, and dataset split strategy on model performance.

Together, these results position WIO-ReefFish not only as a publicly available dataset resource, but also as a more ecologically realistic framework for evaluating computer-vision approaches for coral reef monitoring. As imaging systems and learning approaches continue to evolve, it provides a foundation for the development of more scalable and non-destructive approaches to marine biodiversity assessment.

## CRediT authorship contribution statement

Jules Gerard: Conceptualization, Methodology, Formal analysis, Data curation, Investigation, Writing – original draft, Writing – review and editing.

Luca Branger: Data curation, Investigation, Writing – review and editing.

Filip Huyghe: Investigation, Writing – review and editing.

Marc Kochzius: Conceptualization, Funding acquisition, Project administration, Investigation, Writing – review and editing.

Levy Otwoma: Investigation, Project administration, Writing – review and editing.

Stephen Bergacker: Investigation, Writing – review and editing.

Lode op’t Roodt: Investigation, Writing – review and editing.

Cyrus Rumisha: Investigation, Project administration, Writing – review and editing.

Leandro Di Bella: Methodology, Formal analysis, Writing – review and editing.

## Permits and ethical approval

Underwater video surveys in Kenya and Tanzania were conducted with the support of local institutional partners and under the relevant permissions required for fieldwork at the surveyed sites. Permits and logistical authorisations were facilitated by KMFRI in Kenya and SUA in Tanzania. The dataset was derived exclusively from non-destructive underwater video imagery.

## Data availability

The WIO-ReefFish dataset is publicly available on Zenodo [44].

## Code availability

The benchmarking and evaluation scripts are currently being prepared for public release. Until the code repository is available, scripts can be requested from the corresponding author.

## A. Supplementary Dataset Information

**Table A.1.** Survey sites, sampling dates, and geographic coordinates for underwater video transects collected in Kenya (2023) and Tanzania (2025). Coordinates are given in decimal degrees (°S, °E).

| Country | Year | Date | Location | Lat. ($^{\circ}$S) | Long. ($^{\circ}$E) |
| --- | --- | --- | --- | --- | --- |
| Kenya | 2023 | 09/03 | Mombasa – Shanzu | 3.97695 | 39.76575 |
|  |  | 10/03 | Mombasa – Bamburi Beach | 4.03225 | 39.73275 |
|  |  | 12/03 | Watamu – Moray Reef | 3.37124 | 40.00748 |
|  |  |  | Watamu – Anthias Reef | 3.35505 | 40.02753 |
|  |  | 13/03 | Watamu – Canyon | 3.36214 | 40.01833 |
|  |  | 15/03 | Diani – Milele | 4.35039 | 39.57683 |
|  |  |  | Diani – Galu | 4.25335 | 39.60977 |
|  |  | 16/03 | Diani – Mwanamochi | 4.30641 | 39.60510 |
|  |  | 18/03 | Kisite – Charlie’s Claw | 4.71477 | 39.36800 |
| Tanzania | 2023 | 23/03 | Unguja – Bawe Island | 6.15280 | 39.13437 |
|  |  | 24/03 | Unguja – Murogo Island | 6.16739 | 39.12811 |
|  |  | 27/03 | Chumbe Island – MPA side | 6.27575 | 39.17458 |
| Tanzania | 2025 | 07/02 | Tanga – Fish Eagle Point (Site 1) | 4.84250 | 39.21778 |
|  |  | 08/02 | Tanga – Fish Eagle Point (Site 2) | 4.84972 | 39.21722 |
|  |  | 10/02 | Kunduchi – Bongoyo Island (Site 1) | 6.70278 | 39.28500 |
|  |  |  | Kunduchi – Bongoyo Island (Site 2) | 6.70472 | 39.28694 |
|  |  | 11/02 | Kunduchi – Mbudya MPA (Site 3) | 6.61528 | 39.24500 |
|  |  |  | Kunduchi – Mbudya MPA (Site 4) | 6.61083 | 39.24556 |
|  |  | 14/02 | Kilwa – Somanga (Site 1) | 8.38445 | 39.33565 |
|  |  |  | Kilwa – Somanga (Site 2) | 8.38637 | 39.34095 |
|  |  | 16/02 | Mikindani (Site 1) | 10.24611 | 40.14333 |
|  |  |  | Mikindani (Site 2) | 10.23694 | 40.17222 |
|  |  | 17/02 | Mikindani (Site 3) | 10.23722 | 40.18833 |
|  |  |  | Mikindani (Site 4) | 10.23778 | 40.18750 |
|  |  | 21/02 | Chumbe Island – non-MPA side | 6.27889 | 39.17417 |
|  |  | 22/02 | Chumbe Island – MPA side | 6.28070 | 39.17940 |
|  |  | 25/02 | Nungwi – Tumbatu Island (Site 1) | 5.79194 | 39.21750 |
|  |  | 26/02 | Nungwi – Tumbatu Island (Site 2) | 5.76139 | 39.21722 |
|  |  |  | Nungwi – Tumbatu Island (Site 3) | 5.73944 | 39.28806 |

**Table A.2.** Difficulty-level definitions based on ground-truth bounding-box height percentiles.

| Level | Min. Height | Threshold | GT Subset |
| --- | --- | --- | --- |
| Easy | $h_{55}$ | $\geq 64$ px | $\sim 46\%$ of GT |
| Moderate | $h_{20}$ | $\geq 32$ px | $\sim 80\%$ of GT |
| Hard | — | $\geq 1$ px | 100% of GT |

## B. Implementation Details

### B.1. Training Hyperparameters

YOLO11, YOLOv8, YOLO26, YOLO-World and RT-DETR were trained using AdamW with an initial learning rate of 10^−3^ and cosine learning-rate scheduling. Faster R-CNN, RetinaNet and DINOv2-FRCNN used SGD (*lr* = 10^−2^, momentum = 0.9, weight decay = 5 ×10^−4^) with step-based scheduling. Standard Ultralytics augmentations were applied, including mosaic augmentation, random affine transformations, HSV jitter and horizontal flipping.

Non-maximum suppression (NMS) was applied at IoU = 0.50. All detectors used a confidence threshold of *τ* = 0.25, except Grounding DINO (*τ* = 0.30) to account for noisier zero-shot confidence scores.

Approximate wall-clock training time ranged from ∼0.3 h for YOLO11/YOLOv8 to ∼3.5 h for RT-DETR and Grounding DINO fine-tuning. The full benchmark required fewer than 30 GPU-hours on a single NVIDIA RTX 4090 GPU.

### B.2. Open-Vocabulary Models

Grounding DINO was evaluated in zero-shot mode using a period-separated text prompt formed by concatenating all WIO-ReefFish taxonomic categories (e.g. “`Acanthuridae. Balistidae. Caesionidae. …. Zanclidae`.”). No fine-tuning was performed.

YOLO-World was fine-tuned on the WIO-ReefFish training split using the same category prompts while freezing the text encoder and updating the visual backbone.

### B.3. DINOv2 Detection Head

DINOv2 (ViT-S/14) was evaluated using a lightweight FCOS-style anchor-free detection head trained on frozen backbone features. The adaptation consisted of a 1×1 convolutional feature projection followed by a three-level feature pyramid network constructed from patch tokens up-sampled to multiple spatial resolutions. Separate classification and box-regression branches were used for detection. The detection head was trained for 60 epochs using AdamW with learning rate 10^−3^ and batch size 8.

## C. Taxonomic Performance Analyses

### C.1. Per-class detection performance

Table C.1 reports per-class precision, recall, F1, AP_50_ and AP_50-95_ for RT-DETR under class-aware evaluation. Low-support taxa should be interpreted cautiously.

**Table C.1.** Per-class detection performance of RT-DETR under class-aware evaluation on the test split. Taxa are ordered by test-set support (*n_GT_*). AP values are reported as means with subscripted standard deviations; precision, recall, and F1 correspond to the seed-0 run.

| **Taxon** | $n_{GT}$ | **P** | **R** | **F1** | **$AP_{50}$** | **$AP_{50-95}$** |
| --- | --- | --- | --- | --- | --- | --- |
| Pomacentridae | 470 | .400 | .789 | .531 | .623<sub>.014</sub> | .371<sub>.002</sub> |
| Acanthuridae | 135 | .520 | .659 | .582 | .570<sub>.026</sub> | .432<sub>.018</sub> |
| Holocentridae | 84 | .570 | .869 | .689 | .777<sub>.020</sub> | .556<sub>.005</sub> |
| Labridae | 81 | .400 | .420 | .410 | .343<sub>.030</sub> | .266<sub>.012</sub> |
| Pempheridae | 56 | .639 | .821 | .719 | .694<sub>.045</sub> | .472<sub>.022</sub> |
| Scarinae | 37 | .476 | .541 | .506 | .377<sub>.025</sub> | .310<sub>.026</sub> |
| Balistidae | 25 | .417 | .800 | .548 | .748<sub>.061</sub> | .634<sub>.044</sub> |
| Chaetodontidae | 23 | .565 | .565 | .565 | .566<sub>.026</sub> | .392<sub>.014</sub> |
| Lutjanidae | 14 | .316 | .429 | .364 | .370<sub>.025</sub> | .242<sub>.005</sub> |
| Zanclidae | 12 | .615 | .667 | .640 | .769<sub>.138</sub> | .515<sub>.043</sub> |
| *Taxa with $n_{GT} \leq 10$* |  |  |  |  |  |  |
| Microdesmidae | 10 | .375 | .300 | .333 | .372<sub>.128</sub> | .259<sub>.095</sub> |
| Siganidae | 10 | .250 | .300 | .273 | .356<sub>.041</sub> | .302<sub>.043</sub> |
| Monacanthidae | 9 | .000 | .000 | .000 | .030<sub>.043</sub> | .021<sub>.030</sub> |
| Mullidae | 9 | .500 | .667 | .571 | .472<sub>.066</sub> | .219<sub>.032</sub> |
| Anthiidae | 8 | .750 | .750 | .750 | .567<sub>.100</sub> | .366<sub>.093</sub> |
| Priacanthidae | 7 | .240 | .857 | .375 | .531<sub>.076</sub> | .412<sub>.042</sub> |
| Haemulidae | 5 | .667 | .800 | .727 | .927<sub>.103</sub> | .589<sub>.065</sub> |
| Aulostomidae | 4 | .500 | .500 | .500 | .424<sub>.113</sub> | .326<sub>.147</sub> |
| Caesionidae | 4 | .800 | 1.00 | .889 | .909<sub>.129</sub> | .661<sub>.111</sub> |
| Epinephelidae | 4 | .100 | .250 | .143 | .136<sub>.111</sub> | .091<sub>.072</sub> |
| Pomacanthidae | 3 | .333 | .333 | .333 | .364<sub>.000</sub> | .364<sub>.000</sub> |
| Tetraodontidae | 3 | .250 | .333 | .286 | .098<sub>.033</sub> | .076<sub>.026</sub> |
| Kyphosidae | 1 | .000 | .000 | .000 | .500<sub>.408</sub> | .417<sub>.327</sub> |
| Lethrinidae | 1 | .000 | .000 | .000 | .000<sub>.000</sub> | .000<sub>.000</sub> |
| **Frequent only** | 928 | .443 | .726 | .551 | .540<sub>.011</sub> | .380<sub>.004</sub> |
| **All 24 classes** | 1015 | .443 | .726 | .551 | .480<sub>.015</sub> | .346<sub>.002</sub> |

### C.2. Confusion matrices

Confusion matrices for the two strongest-performing supervised detectors, RT-DETR and YOLO11, are shown in Figs. C.2 and C.3, respectively.

**Table C.2.**
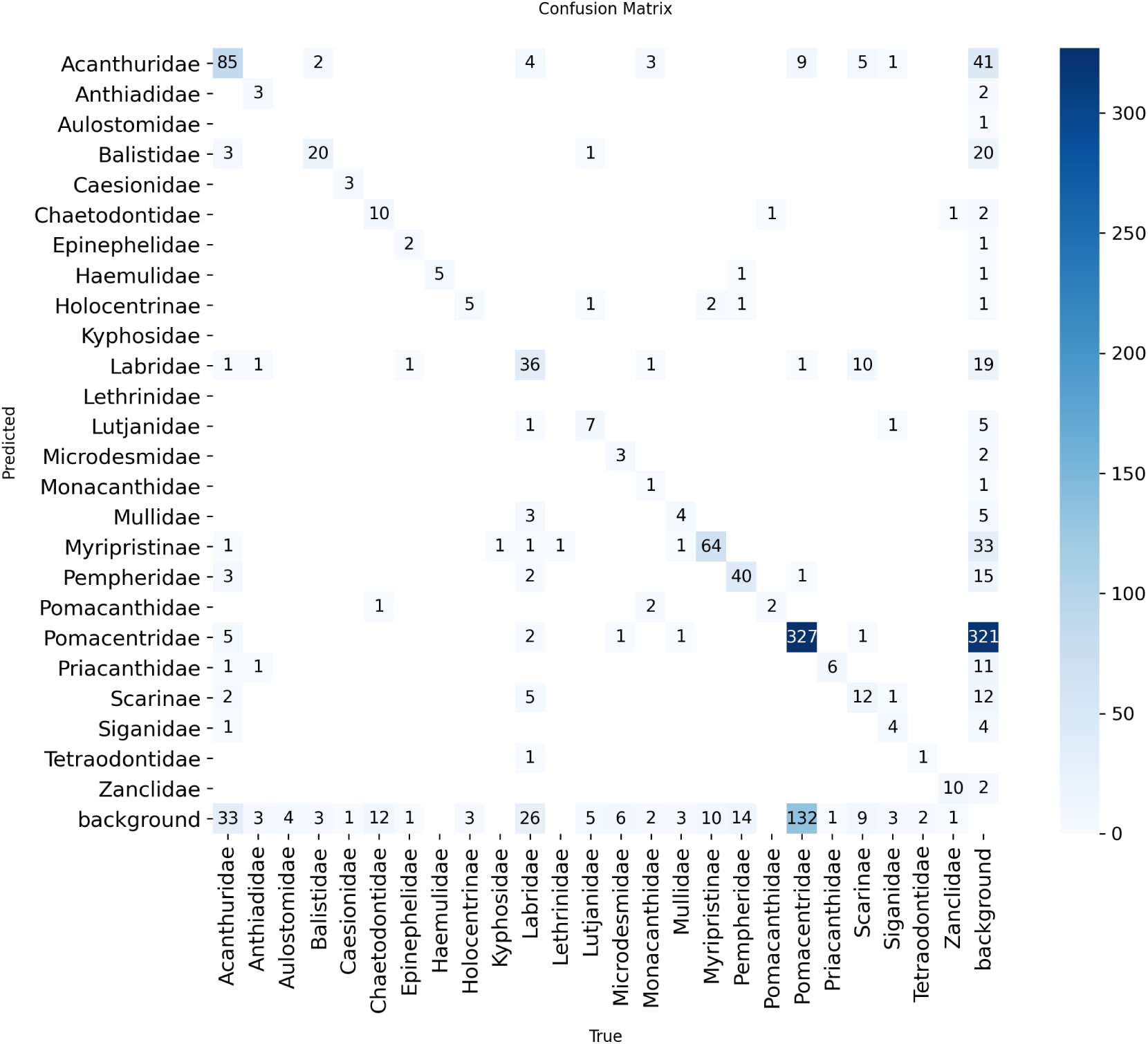
Confusion matrix for RT-DETR under the class-aware protocol on the test split. Rows indicate predicted labels and columns indicate true labels. Values are shown as absolute counts.

**Table C.3.**
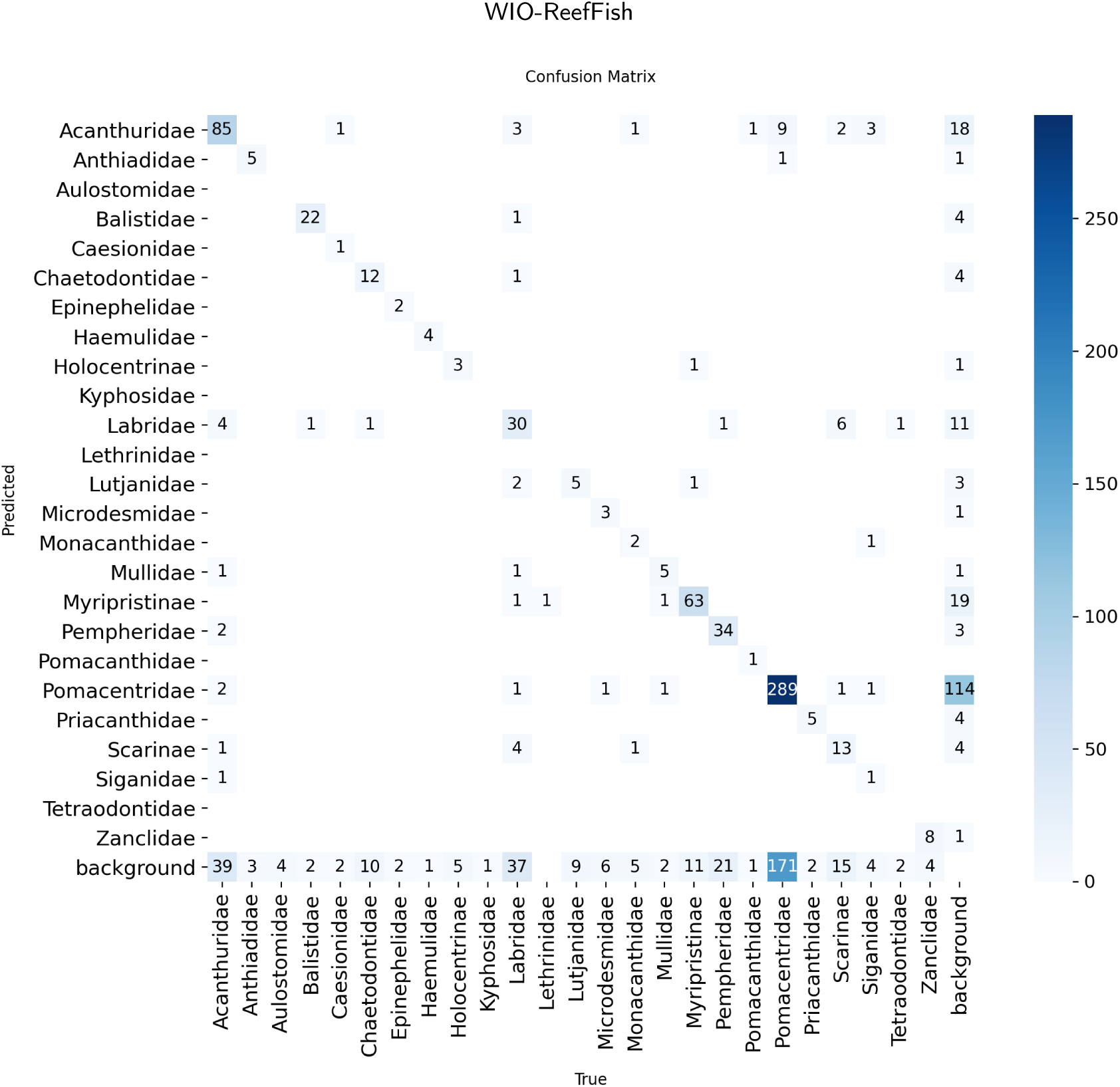
Confusion matrix for YOLO11 under the class-aware protocol on the test split. Rows indicate predicted labels and columns indicate true labels. Values are shown as absolute counts.

## D. Additional Detection Analyses

### D.1. Per-location performance

**Table D.1.**
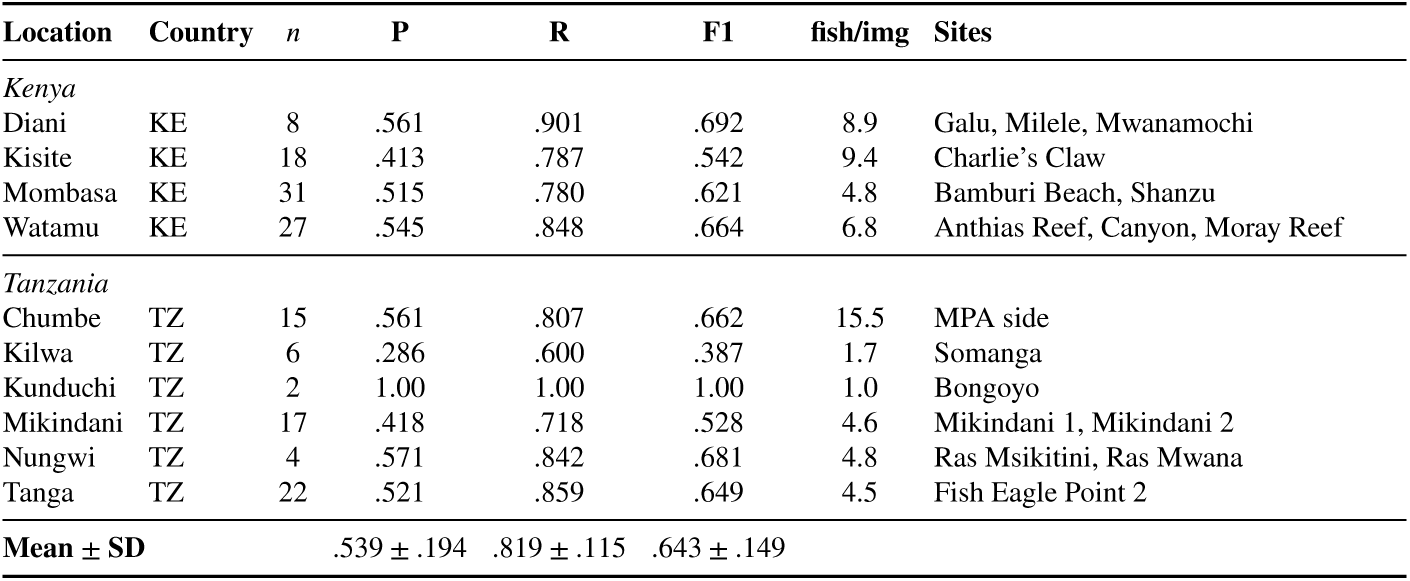
Per-location detection performance of RT-DETR under class-agnostic evaluation on the test split. Results are aggregated over all sites within each location. *n* denotes the number of test images, and *fish/img* the mean annotation density.

### D.2. Difficulty-stratified performance

**Table D.2.**
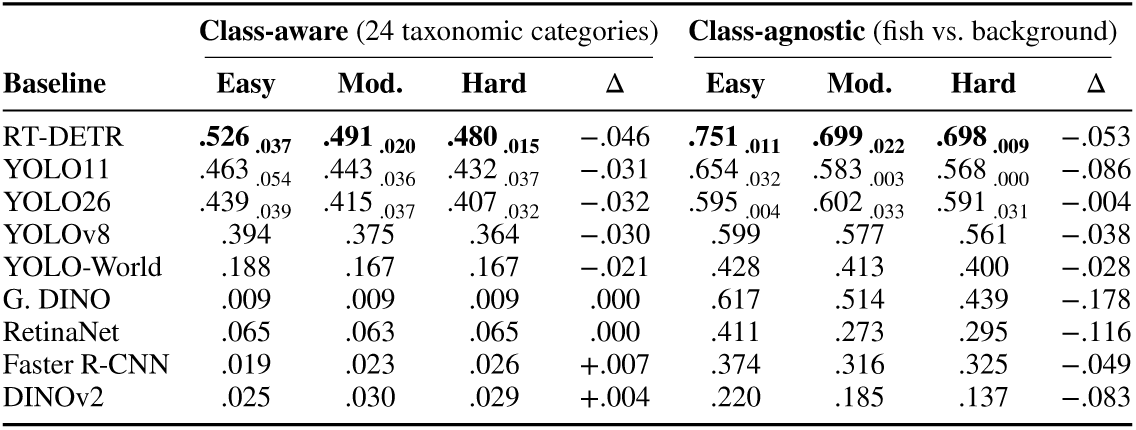
mAP_50_ by difficulty level under class-aware and class-agnostic evaluation on the test split. Δ denotes the change from Easy to Hard. Subscripted values denote standard deviations.

### D.3. Brightness-stratified performance

Fish detections were stratified into dark, moderate and bright categories based on HSV brightness values computed within each annotated bounding box. Thresholds were derived from the 25th and 75th percentiles of brightness values across the test set.

**Table D.3.** F1 score by ground-truth brightness level under the class-agnostic protocol on the test split. Brightness groups correspond to quartile-based HSV V-channel thresholds computed from test-set annotations.

| Baseline | Dark ($n=254$) | Moderate ($n=507$) | Bright ($n=254$) |
| --- | --- | --- | --- |
| RT-DETR | **.736** | **.826** | **.854** |
| YOLO26 | .602 | .643 | .713 |
| YOLO11 | .587 | .653 | .724 |
| YOLOv8 | .579 | .633 | .724 |
| G. DINO | .508 | .554 | .685 |
| RetinaNet | .512 | .600 | .602 |
| Faster R-CNN | .492 | .629 | .594 |
| YOLO-World | .425 | .489 | .386 |
| DINOv2 | .201 | .168 | .256 |

All detectors showed progressively higher F1 scores from dark to bright fish. RT-DETR remained the strongest-performing detector across all brightness categories, increasing from 0.736 F1 for dark fish to 0.854 for bright fish. Background brightness and luminance contrast showed only weak associations with location-level F1 (*r* = 0.07 and *r* = −0.25, respectively).

### D.4. Weighted Box Fusion

Weighted Box Fusion (WBF) [45] was evaluated using the five strongest-performing detectors under the class-agnostic protocol. Lower-performing detectors were excluded from fusion experiments because preliminary tests showed that their high false-positive rates degraded ensemble performance.

**Table D.4.** Weighted Box Fusion results under the class-agnostic protocol on the test split. Fusion was restricted to the five strongest-performing detectors: RT-DETR, YOLO11, YOLO26, YOLOv8 and YOLO-World.

| Model | P | R | F1 | TP | FP | FN |
| --- | --- | --- | --- | --- | --- | --- |
| YOLO11 | .659 | .654 | .656 | 664 | 344 | 351 |
| YOLO26 | .658 | .650 | .654 | 660 | 343 | 355 |
| YOLOv8 | .636 | .642 | .639 | 652 | 373 | 363 |
| RT-DETR | .504 | **.811** | .622 | 823 | 809 | 192 |
| YOLO-World | **.674** | .447 | .538 | 454 | 220 | 561 |
| **ENSEMBLE (WBF, top-5)** | **.775** | .694 | **.732** | 704 | 204 | 311 |

WBF improved F1 by 0.076 relative to the best individual 640 × 640 model, mainly by increasing precision while retaining moderate recall.

### D.5. Inference-time image enhancement

Three common underwater image enhancement methods were applied during inference to detectors trained on raw underwater imagery.

**Table D.5.** Effect of underwater image enhancement on class-aware F1 on the test split. Δ is the F1 change relative to raw, unenhanced inference for the same baseline.

| Baseline | Raw | CLAHE | White Bal. | Sea-thru |
| --- | --- | --- | --- | --- |
| YOLO26 | **.585** | .468 $\downarrow_{.117}$ | .542 $\downarrow_{.043}$ | .442 $\downarrow_{.143}$ |
| YOLO11 | .571 | .404 $\downarrow_{.167}$ | .480 $\downarrow_{.091}$ | .390 $\downarrow_{.182}$ |
| YOLOv8 | .566 | .421 $\downarrow_{.145}$ | **.537** $\downarrow_{.028}$ | .439 $\downarrow_{.127}$ |
| RT-DETR | .542 | .426 $\downarrow_{.116}$ | .519 $\downarrow_{.023}$ | .466 $\downarrow_{.076}$ |
| YOLO-World | .142 | .069 $\downarrow_{.073}$ | .133 $\downarrow_{.009}$ | .095 $\downarrow_{.047}$ |
| **Avg. ΔF1** | — | −.124 | −.039 | −.115 |

All enhancement methods reduced detection performance. White balance had the smallest average effect (−0.039), while CLAHE and Sea-thru produced larger declines (−0.124 and −0.115, respectively). This pattern is consistent with a train/test photometric mismatch rather than an intrinsic limitation of the enhancement methods.

## Acknowledgements

We thank the field teams, divers, local collaborators, partner institutions and authorities who supported underwater video data collection, field logistics and site access in Kenya and Tanzania. We acknowledge the institutional support of Vrije Universiteit Brussel (VUB), the Kenya Marine and Fisheries Research Institute (KMFRI), and Sokoine University of Agriculture (SUA).

